# Tunable Transcriptional Interference at the Endogenous Alcohol Dehydrogenase Gene Locus in *Drosophila melanogaster*

**DOI:** 10.1101/452649

**Authors:** Victoria Jorgensen, Jingxun Chen, Helen Vander Wende, Devon Harris, Siu Wah Wong-Deyrup, Yuzhang Chen, Prashanth Rangan, Gloria Ann Brar, Eric M. Sawyer, Leon Chan, Elçin Ünal

**Author notes:** These authors contributed equally to this work (Victoria Jorgensen, Jingxun Chen, Helen Vander Wende).

## Abstract

Neighboring sequences of a gene can influence its expression. In the phenomenon known as transcriptional interference, transcription at one region in the genome can repress transcription at a nearby region in *cis*. Transcriptional interference occurs at a number of eukaryotic loci, including the alcohol dehydrogenase (*Adh*) gene in *Drosophila melanogaster. Adh* is regulated by two promoters, which are distinct in their developmental timing of activation. It has been shown using transgene insertion that when the promoter distal from the *Adh* start codon is deleted, transcription from the proximal promoter becomes de-regulated. As a result, the *Adh* proximal promoter, which is normally active only during the early larval stages, becomes abnormally activated in adults. Whether this type of regulation occurs in the endogenous *Adh* context, however, remains unclear. Here, we employed the CRISPR/Cas9 system to edit the endogenous *Adh* locus and found that removal of the distal promoter does also result in the untimely expression of the proximal promoter-driven mRNA isoform in adults, albeit at lower levels than previously reported. Importantly, we show that transcription from the distal promoter is sufficient to repress proximal transcription in larvae and that the degree of this repression depends on the degree of distal promoter activity. Finally, repression of the endogenous *Adh* proximal promoter is associated with the enrichment of histone 3 lysine 36 trimethylation (H3K36me3), a chromatin mark necessary for transcription-coupled gene repression in yeast. We conclude that the endogenous *Adh* locus is developmentally regulated by transcriptional interference in a tunable manner.

## Introduction

Transcriptional interference, or *cis*-mediated downregulation of transcription at a locus as a result of transcription from a nearby location (Shearwin et al., 2005), was initially recognized as a mechanism of gene regulation conferred by retroviral promoters (Cullen et al., 1984). Since then, transcriptional interference has been observed to endogenously regulate genes in a number of eukaryotic contexts (Martens et al., 2004; Bird et al., 2006; Hongay et al., 2006; Hainer et al., 2011; van Werven et al., 2012; reviewed in Shearwin et al., 2005). In particular, transcription of non-coding RNAs is widely associated with interference of promoters or regulatory elements of local coding transcripts (Martens et al., 2004; Hongay et al., 2006; van Werven et al., 2012; Kaikkonen and Adelman, 2018). In addition to non-coding RNAs, mRNA isoforms have also been linked to transcriptional interference. For genes with more than one promoter, transcription from the distal promoter may not only produce a distinct mRNA isoform, but could also lead to the repression of an mRNA isoform transcribed from the open reading frame (ORF)-proximal gene promoter (Corbin and Maniatis, 1989; Moseley et al., 2002; Sehgal et al., 2008; Liu et al., 2015; Chen et al., 2017). In addition, since distinct mRNA isoforms may differ in their translational efficiency, regulation of promoter choice may impact gene expression at the protein level. In some instances, this difference in translational efficiency is due to the presence of upstream ORFs (uORFs) in the 5’ leader of the distal promoter-derived mRNA isoform, which could inhibit translation of the protein-coding ORF (Moseley et al., 2002; Law et al., 2005; Sehgal et al., 2008; Ingolia et al., 2011; Rojas-Duran and Gilbert, 2012; Brar et al., 2012; Chew et al., 2016; Bird and Labbé, 2017; Chen et al., 2017; Cheng et al., 2018; Zhang et al., 2018). As a result, in these cases, transcription of a distal promoter-derived mRNA isoform causes downregulation of protein expression through the integration of two seemingly disparate mechanisms of transcriptional and translational repression (Chen et al., 2017; Cheng et al., 2018; Van Dalfsen et al., 2018; Hollerer et al. submitted).

Transcription can antagonize downstream promoter activity by at least two means: First, the movement of the transcription machinery through the downstream promoter could interfere with transcription factor binding (reviewed in Shearwin et al., 2005; van Werven et al., 2012; Zafar et al., 2014; Chia et al., 2017). Second, transcription through the downstream promoter could establish a repressive chromatin state (Hainer et al., 2011; van Werven et al., 2012; Woo et al., 2017; Chia et al., 2017). These mechanisms are not mutually exclusive and in fact have been shown to act in concert (van Werven et al., 2012; Chia et al., 2017). In the case of chromatin state changes, co-transcriptional histone modifications such as histone 3 lysine 36 trimethylation (H3K36me3) have been associated with nucleosome stabilization and repression of the downstream promoter (Hampsey and Reinberg, 2003; Carrozza et al., 2005; Keogh et al., 2005; Houseley et al., 2008; Kim and Buratowski, 2009; Govind et al., 2010; Kim et al., 2012; van Werven et al., 2012; Ard and Allshire, 2016; Chia et al., 2017). While these studies have been conducted extensively in yeast, the link between H3K36me3 and transcription-coupled repression has been less clear in metazoans.

An established example of transcriptional interference is the regulation of the alcohol dehydrogenase (*Adh*) gene in the fruit fly, *Drosophila melanogaster* (Corbin and Maniatis 1989). *Adh* is transcribed from two closely positioned promoters, resulting in the production of at least two distinct mRNA isoforms (Figure 1A). These transcript isoforms are expressed in a developmentally regulated and tissue-specific manner (Ursprung et al., 1970; Benyajati et al., 1983; Savakis et al. 1986; Anderson et al., 1991; Visa et al., 1992; reviewed in Sofer and Martin, 1987). Transcription occurs from the ORF-proximal promoter (hereon referred to as *Adh* proximal promoter) during the early larval stages and from the ORF-distal promoter (hereon referred to as *Adh* distal promoter) during late third instar larvae and in adults (Fig. 1B, adapted from Corbin and Maniatis, 1989 as well as Sofer and Martin, 1987). It has been shown that transcription from the *Adh* distal promoter is necessary to repress transcription from the *Adh* proximal promoter (Corbin and Maniatis, 1989). However, this previous study employed transgene insertions, and the same allele displayed variable degrees of transcriptional interference, attributed to positional effects (Corbin and Maniatis, 1989). Therefore, both the impact and the extent of transcriptional interference at the endogenous *Adh* locus are currently unknown. It also remains to be tested whether the premature expression of the *Adh* distal transcript in larvae is sufficient to down-regulate the *Adh* proximal promoter. Furthermore, whether transcription from the *Adh* distal promoter is accompanied by downstream changes in H3K36me3 is unknown. Finally, the translational capacity of the two *Adh* mRNA isoforms has not been investigated. Here, we examined these unexplored aspects of *Drosophila Adh* regulation. We report that the transcriptional interference at the endogenous *Adh* locus is tunable and associated with H3K36me3 enrichment. We further show that the two *Adh* transcript isoforms are both associated with high polysome fractions, indicating efficient translation.

**Fig. 1.**
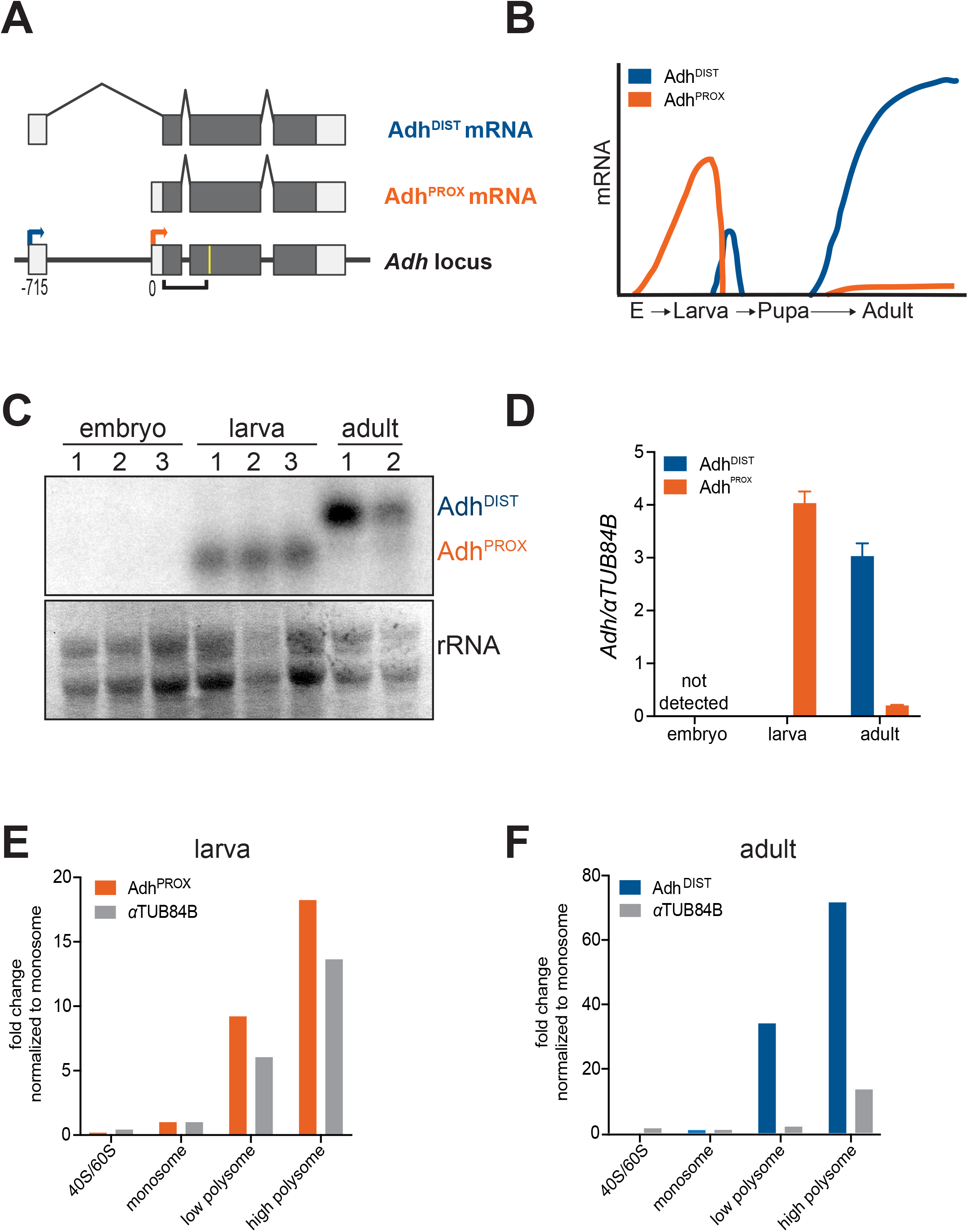
**Transcription and translation of the two *Adh* isoforms during *Drosophila* development. (A)** Illustration of coding (grey) and non-coding (white) exons of the *Adh* locus and the two *Adh* mRNA isoforms. Transcription of *Adh* can occur at either of two distinct transcription start sites (TSSs): the proximal TSS (orange arrow), nearest to the gene body, produces a short mRNA transcript (Adh^PROX^), while the distal TSS (blue arrow), farthest from the gene body, produces a 5’ extended mRNA (Adh^DIST^). Numbers below the *Adh* locus refer to distance in base pairs (bp) from the Adh^PROX^ TSS. The yellow line represents the relative location of the oligonucleotide probe used for RNaseH cleavage and the black-bracketed line represents the probe used in RNA blotting shown in **(C). (B)** Schematic adapted from (Corbin and Maniatis, 1989) and (Sofer and Martin, 1987) showing expression of *Adh* mRNA isoforms throughout development. **(C)** RNA blot of wild-type *Drosophila* RNA extracts throughout development confirms the stage-specific expression of both isoforms. Embryos were collected at 8 hours and were collected at 72 hours, corresponding. *Adh* transcripts were detected using a probe that hybridizes to a common region of all isoforms. Because the two isoforms vary by only ~50 bp, all samples were RNaseH cleaved in the second exon for better separation. Methylene blue staining of rRNA was used as a loading control. Numbered lanes indicate biological replicates. **(D)** Expression levels of Adh^PROX^ and Adh^DIST^ measured by RT-qPCR using isoform-specific primers. All data were normalized to *αTUB84B*. The mean of two independent repeats from two separate collections is shown. Error bars represent the range. **(E)** and **(F)** RT-qPCR analysis of polysome profiles for Adh^PROX^ (orange), Adh^DIST^ (blue) and a control αTUB84B transcript (black) in wild-type L1/L2 larvae harvested at 80 hours **(E)** and wild-type adults **(F)**. RNA was isolated individually from fractions and pooled into four categories: 40S/60S, monosome, low polysome (di- and trisome), high polysome (remaining fractions). Expression levels were obtained using isoform-specific primers and RT-qPCR. Data were first normalized to *in vitro* transcribed *RCC1*, which was spiked at equal amounts into each fraction prior to RNA extraction. Normalized data were then plotted relative to the amount present in the monosome fraction for each transcript.

## Results

The *Adh* proximal promoter produces a transcript of 1001 nucleotides in length (hereon referred to as Adh^PROX^), whereas the *Adh* distal promoter activates a transcription start site (TSS) located 715 base pairs (bps) upstream of the proximal TSS. The resulting transcript from the distal promoter, hereon referred to as Adh^DIST^, has a unique 5’ leader located in exon 1 (Figure 1A). We first measured the relative abundance of the two *Adh* mRNA isoforms from wild-type embryos, larvae, and adult fruit flies using RNA blot hybridization. Because the two *Adh* isoforms differ by only 56 nucleotides, we employed an RNaseH digestion strategy to shorten the full-length transcripts so that a clear difference in isoform length could be detected (Figure 1A). Consistent with previous work (Savakis et al., 1986; Corbin and Maniatis, 1989; diagrammed in Figure 1B), we observed that both *Adh* transcripts were undetectable in embryos (Figure 1C). Adh^PROX^ specifically was expressed at high levels in early larval stages, and the Adh^DIST^ transcript was the predominant isoform in adults. To quantitate the relative expression levels of each isoform, we used reverse transcription followed by quantitative Polymerase Chain Reaction (RT-qPCR) using isoform-specific primers. This analysis revealed that the Adh^PROX^ transcript level was ~20 fold higher in larvae compared to in adults, whereas the Adh^DIST^ transcript had the reciprocal pattern with more than 8000-fold enrichment in adults compared to its expression level in larvae. (Figure 1D). These data confirm that the *Adh* locus undergoes dramatic, developmentally induced transcript isoform toggling, as evidenced by the mutually exclusive expression patterns of the two mRNA isoforms.

To determine the translational status of the two *Adh* isoforms, we enriched for ribosome-associated transcripts using sucrose gradient fractionation and measured the relative distribution of Adh^PROX^ or Adh^DIST^ across different fractions in larvae and whole adults. Adh^PROX^ was enriched in the polysome fractions to a similar extent compared to a well-translated, ubiquitous transcript, αTUB84B (Figure 1E and Figure S1A). Interestingly, Adh^DIST^ enrichment in the polysome fractions from adults was more than 8-fold higher relative to αTUB84B enrichment (Figure 1F and Figure S1B). We conclude that both Adh^PROX^ and Adh^DIST^ are well translated. Furthermore, Adh^DIST^ appears to be noticeably more enriched in the polysome fractions than either Adh^PROX^ or αTUB84B, indicating enhanced translational efficiency.

To assess the impact of transcriptional interference on Adh^PROX^ expression at the endogenous locus, we used CRISPR/Cas9-based editing to delete the *Adh* distal promoter (*Adh^DIST∆^*) (Figure 2A and Figure 2B). Deletion of the *Adh* distal promoter resulted in a dramatic reduction of the Adh^DIST^ transcript and led to the expression of Adh^PROX^ in both larvae and adults, albeit at low levels (Figure 2C and Figure S2). RT-qPCR analysis showed a 6-fold increase in Adh^PROX^ abundance in *Adh^DIST∆^* mutants compared to wild-type adults (Figure 2D). We conclude that loss of transcription from the *Adh* distal promoter results in a modest de-repression of Adh^PROX^, suggesting that, at least in adult fruit flies, transcription from the *Adh* distal promoter antagonizes the activity of the *Adh* proximal promoter.

**Fig. 2.**
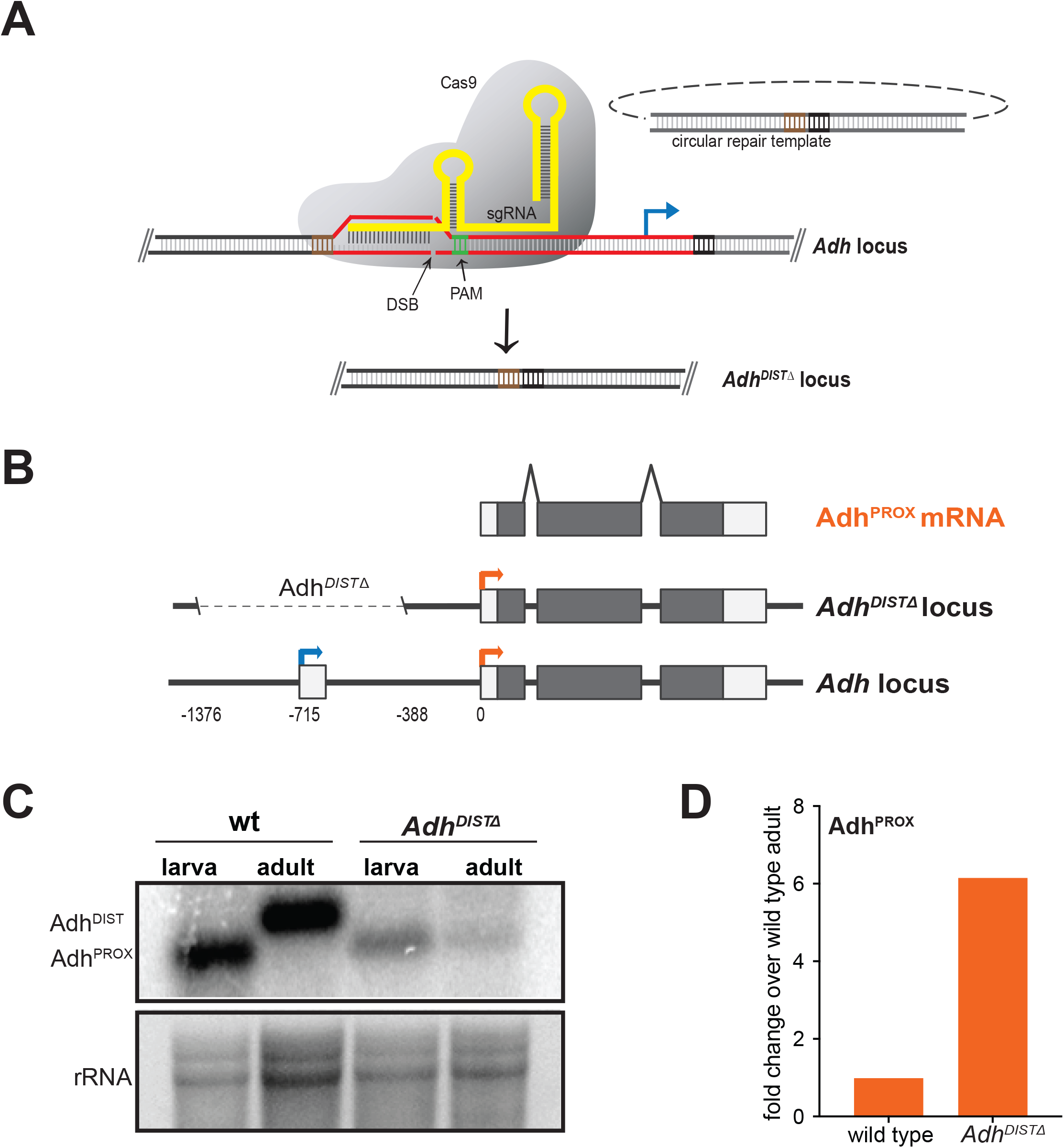
**Deletion of the endogenous *Adh^DIST^* promoter leads to Adh^PROX^ expression in adults. (A)** Schematic depicting CRISPR-Cas9 editing scheme to delete the *Adh^DIST^* promoter region. A double-strand break (DSB) was made at the PAM site (green) using Cas9 nuclease (grey) and short guide RNA (sgRNA) (yellow), which binds ~730 base pairs (bp) upstream of the Adh^DIST^ transcription start site. The DNA region in red signifies the region that was deleted. The DNA region in black and brown signifies the DNA sequence flanking the deleted region. The blue arrow refers to the transcription start site of Adh^DIST^. Figure is not drawn to scale. **(B)** Schematic of *Adh^DIST^* promoter deletion, which will be referred to as *Adh^DIST∆^*. Coding (grey) and non-coding (white) exons are shown. Arrows represent TSS of Adh^PROX^ (orange) and Adh^DIST^ (blue). Numbers below the *Adh* locus refer to distance in base pairs (bp) from the Adh^PROX^ TSS. **(C)** RNA blot in wild-type and *Adh^DIST∆^* adult fruit flies and L1/L2 larvae. RNA isoforms were detected using a probe that hybridizes to a common region of all isoforms. Methylene blue staining of rRNA was used as a loading control. **(D)** Expression levels of Adh^PROX^ measured by RT-qPCR using isoform-specific primers. Data were first normalized to *αTUB84B* and then to wild-type adult levels and are shown as fold change over wild type.

Next, we tested if untimely overexpression of Adh^DIST^ during larval development was sufficient to repress Adh^PROX^ expression. Employing a similar CRISPR/Cas9-based editing strategy, we replaced the endogenous *Adh* distal promoter with an inducible *10xUAS*-*hsp70* promoter (*Adh^UAS^*, transcript produced from this promoter is also referred to as Adh^DIST^ for simplicity) (Figure 3A). The *Adh^UAS^* line was crossed to a *tub-GAL4* line, which exhibits ubiquitous *GAL4* expression driven from the *αTub84B* promoter. In the F_1_ larvae, we observed ~3000-fold increase of the Adh^DIST^ isoform compared to wild type, accompanied by ~10-fold decrease in the Adh^PROX^ isoform (Figure 3B and 3C). We noticed that, in F1 larvae from the *Adh^UAS^* lines, Adh^DIST^ expression was apparent even without the *GAL4* driver, likely due to leaky expression from the *hsp70* promoter, located immediately upstream of the Adh^DIST^ TSS (Figure 3C and Figure S3). Comparison of lines with and without GAL4 thus allowed us to achieve a range of Adh^DIST^ expression levels, which provided insight into the dose-dependent relationship between production of Adh^DIST^ and Adh^PROX^. We found that the degree of Adh^DIST^ overexpression scaled with the degree of Adh^PROX^ repression: the more the distal promoter activity, the less the proximal transcript abundance (Figure 3C). This observation suggests that the antagonistic relationship between the levels of the two transcript isoforms is not binary, but tunable. RNA blotting confirmed that Adh^DIST^ levels were highest in lines carrying the GAL4 driver. Adh^DIST^ was also expressed in *Adh^UAS^* homozygous lines without the GAL4 driver (Figure 3B). In *Adh^UAS^* heterozygous lines without the GAL4 driver, Adh^DIST^ expression in F1 larvae was barely detectable by RNA blotting, but still higher than wild-type larvae, consistent with the RT-qPCR data (Figure 3B and 3C). We conclude that Adh^DIST^ transcription is sufficient to downregulate Adh^PROX^ expression in a dose-dependent manner.

**Fig. 3.**
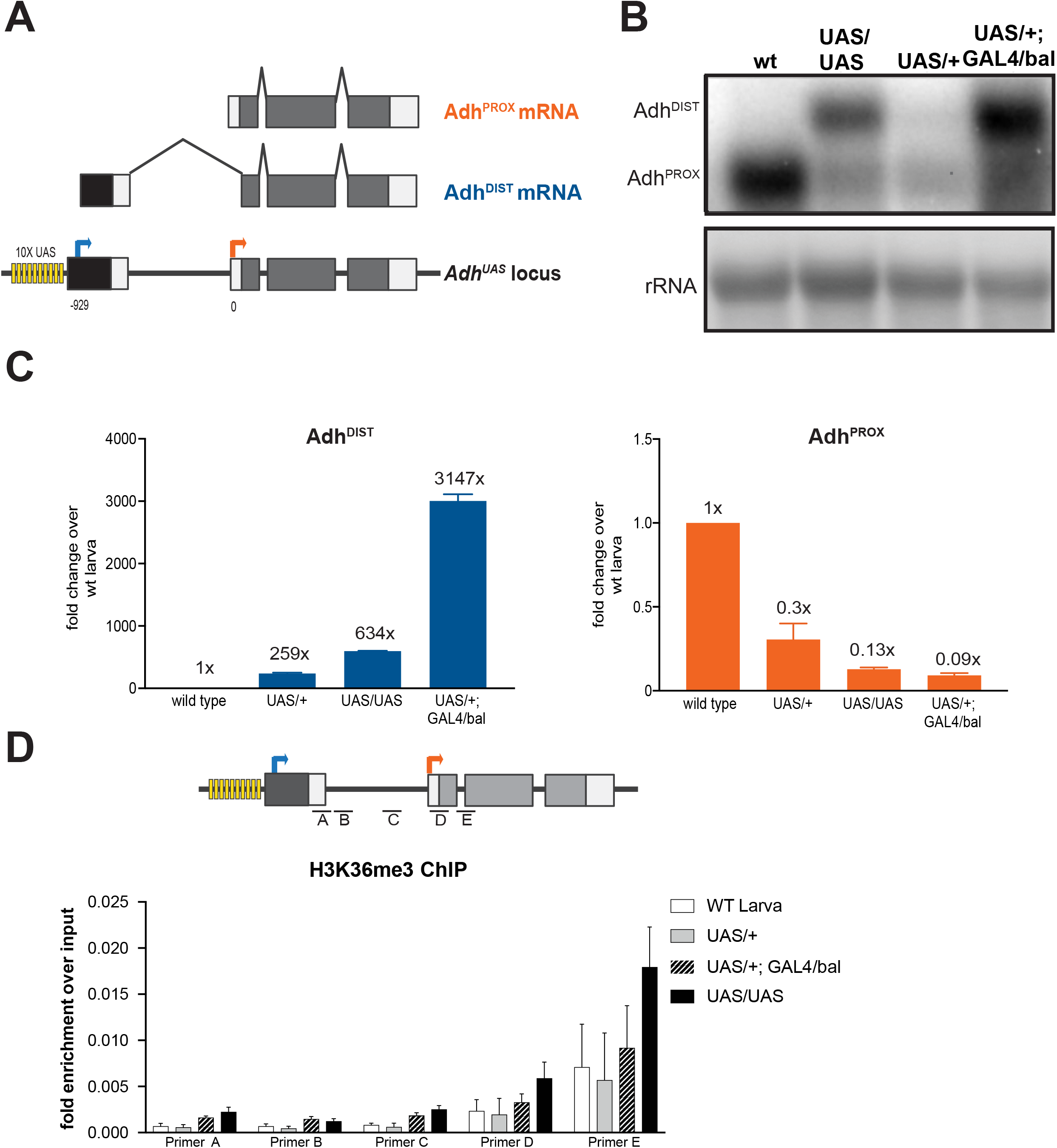
**Ectopic expression of Adh^DIST^ is sufficient for downregulation of Adh^PROX^ in larvae. (A)** Diagram of GAL4/UAS induction system for *Adh*. Immediately upstream of the Adh^DIST^ TSS are 10 consecutive GAL4 bind sites (UAS) (shown in yellow) followed by the minimal *hsp70* promoter (black). Coding (grey) and non-coding (white) exons are shown. Arrows represent TSSs of Adh^PROX^ (orange) and Adh^DIST^ (blue). Numbers below the *Adh* locus refer to distance in base pairs (bp) from the Adh^PROX^ TSS. The TSS for the GAL4-induced isoform (Adh^UAS^, referred to as Adh^DIST^ elsewhere for consistency) was determined by 5’ RACE (Figure supplement 3A). **(B)** RNA blot analysis confirms that ectopic expression of Adh^DIST^ in larvae is sufficient for Adh^PROX^ downregulation. RNA isoforms were detected using a probe that hybridizes to a common region of all isoforms. Methylene blue staining of rRNA was used as a loading control. **(C)** Expression levels of Adh^DIST^ and Adh^PROX^ in larvae with varying degrees of Adh^DIST^ induction. Abundances of Adh^DIST^ (left) and Adh^PROX^ (right) were measured in the following four larval lines: wild-type larva, heterozygous UAS (UAS/+), homozygous UAS (UAS/UAS), heterozygous GAL4 and UAS (UAS/+, GAL4/+). Expression levels were obtained using isoform specific primers and RT-qPCR. All data were normalized to *αTUB84B* and then to wild type (1x). Error bars represent range (n=2 biological replicates) **(D)** Induction of distal transcription promotes histone H3 lysine 36 trimethylation (H3K36me3) over the Adh^PROX^ promoter. DNA recovered from chromatin IP were quantified using RT-qPCR and 5 primer pairs (A, B, C, D, and E) spanning the region between the *Adh* promoters as well as 5’ end of the gene body. The mean of three independent biological repeats and the standard error of the mean are shown.

To test if transcription from the distal promoter led to changes in chromatin marks over the proximal promoter, we performed chromatin immunoprecipitation (ChIP) against H3K36me3 in larvae collected from wild type as well as the three *Adh^UAS^* lines. H3K36me3 is a co-transcriptionally established modification that occurs in regions downstream of active promoters (Xiao et al., 2003; Bannister et al., 2005; Mikkelsen et al., 2007). In yeast, H3K36me3 is associated with a decrease in spurious transcription initiation from within the body of transcribed genes and repression of gene promoters (Hampsey and Reinberg, 2003; Carrozza et al., 2005; Keogh et al., 2005; Houseley et al., 2008; Kim and Buratowski, 2009; Govind et al., 2010; Kim et al., 2012; van Werven et al., 2012; Ard and Allshire, 2016; Chia et al., 2017). We observed an increase in H3K36me3 enrichment downstream of the Adh^DIST^ transcription start site in the two *Adh^UAS^* lines in which the abundance of the Adh^DIST^ transcript and the degree of Adh^PROX^ repression were most pronounced: UAS/UAS and UAS/+; GAL4/bal (Figure 3D). Comparison of these two lines was informative. We observed greater relative enrichment of H3K36me3 over the *Adh* proximal promoter in the former line (Figure 3D), supporting a model in which this chromatin mark was deposited in *cis*, and in a manner that was dependent on Adh^DIST^ transcription. These data are highly reminiscent of what has been reported at the *NDC80* locus in budding yeast (Chia et al., 2017) and *MDM2* locus in human cells (Hollerer et al., submitted).

## Discussion

The fruit fly *Adh* locus, which encodes alcohol dehydrogenase, is a well-established example of transcriptional interference. At the time that it was originally investigated, however, the locus was studied outside of its natural genomic context, using P element transgenes (Corbin and Maniatis, 1989). Here, we revisit the regulation of this locus, leveraging CRISPR/Cas9-based editing, reverse transcription coupled with quantitative PCR, and chromatin immunoprecipitation to better define the regulation of this important gene. Although Adh^PROX^ is the predominant transcript isoform encoding ADH enzyme during normal larval development, we demonstrate that the engineered induction of the Adh^DIST^ transcript is sufficient to repress Adh^PROX^ expression. Importantly, the degree of the distal promoter activity correlates well with the extent of transcriptional interference. Tunable transcriptional interference was first reported in bacteria (Bordoy et al., 2016; Hao et al., 2016), more recently in yeast (Chia et al., 2017), and potentially in human cells (Hollerer et al., submitted). We also found that the tunable transcriptional interference seen at the *Drosophila Adh* locus is associated with the enrichment of H3K36me3 marks over the *Adh* proximal promoter. Similar findings were observed for the kinetochore gene *NDC80* in budding yeast (further discussed below; Chia et al., 2017) and the oncogene *MDM2* in human cells (Hollerer et al., submitted). All of these studies highlight the notion that gene regulation by transcriptional interference is not binary with an on/off state, but rather can be utilized to tune the expression of regulated mRNAs during developmental gene expression programs.

Although deletion of the *Adh* distal promoter at the endogenous locus de-repressed Adh^PROX^ expression in adult fruit flies, the severity of this phenotype was far less pronounced compared to a previous study (Corbin and Maniatis, 1989). A possible explanation for this difference is that position effects arising from differences in P element transgene insertion sites might alter the levels of transcription interference that were observed. The reduction of the Adh^PROX^ transcript in *Adh^DIST∆^* larvae suggests that the deleted region carries sites for some as yet to be determined positive regulators for Adh^PROX^ expression. Alternatively, the deletion could change the nucleosome positioning in this region, which could impact Adh^PROX^ expression. Regardless of these points, our study demonstrates that at the endogenous *Adh* locus, distal promoter-driven transcriptional interference is necessary for Adh^PROX^ repression.

The regulation of the *Adh* gene described here has some similarities to that found for the *NDC80* gene in budding yeast (Chen et al., 2017; Chia et al., 2017). First, both genes have two promoters that are developmentally regulated, with the distal and proximal promoter encoding two distinct mRNA isoforms. Second, transcriptional interference is similar in both cases: transcription from the distal promoter is necessary and sufficient to repress the expression of the proximal promoter-derived isoform. Concomitant with this interference is the enrichment of repressive H3K36me3 marks over the proximal promoter. While the H3K36me3 enrichment is similar between the cases of *Drosophila Adh* and yeast *NDC80*, we have been unable to assess causality in the current study. H3K36me3 is deposited by Set2, a highly conserved methyltransferase that physically associates with the elongating RNA polymerase II (Xiao et al., 2003). *Set2* is essential for the viability of the fruit fly (Bell et al., 2007). Our attempts to characterize SET2 involvement in *Adh* regulation using RNA interference were unsuccessful, since these lines did not survive to adulthood. This finding precluded us from determining the impact of H3K36me3 on Adh^PROX^ expression.

A key difference between examples of *Adh* and *NDC80* gene regulation is related to the translatability of the distal promoter-derived transcript isoforms. In the case of *NDC80*, the ORF within the distal promoter-derived mRNA is not translated, due to competing translation of multiple upstream uORFs that are located in the 5’ leader of this transcript. The *NDC80* case thus shows an interesting link between transcriptional and translational regulation. In essence, production of the distal promoter-derived transcript results in both transcriptional and translational repression, ultimately together resulting in decreased Ndc80 protein production. By contrast, the Adh^DIST^ transcript isoform is well translated, even better than the highly expressed αTUB84B transcript. The lack of translational repression in Adh^DIST^ is consistent with the absence of an AUG start codon within the 5’ leader of this transcript (Figure S3), thus excluding repressive uORF translation. The difference between the apparent regulation in these two cases is important: poor translation in the case of the 5’ extended *NDC80^LUTI^* isoform and superior translation in the case of Adh^DIST^. It is interesting to note that an earlier study, which examined the consequences of a natural transposon insertion at the *Adh* locus in the fruit fly (Dunn and Laurie, 1995), along with a previous report (Laurie and Stam, 1988), showed that the insertion of a *copia* retrotransposon between the *Adh* adult enhancer and the *Adh* distal promoter leads to an unusually low level of the ADH protein and enzyme activity. The reduction was found to occur as a result of a decrease in the level of the Adh^DIST^ transcript. Surprisingly though, in this case, the Adh^PROX^ transcript levels were proportionally increased in adults (Dunn and Laurie, 1995). Given that the levels of the distal and proximal transcripts remain similar between the wild type and the lines carrying transposon insertion, these data suggest that in the adult fruit flies, Adh^PROX^ might not be as efficiently translated as Adh^DIST^, which is consistent with our polysome analysis. One possibility is that tissue-specific, trans-acting factors could differentially modulate the translation of the two *Adh* mRNA isoforms. Such spatial effects are likely to be missed by the whole organism polysome fractionation approach that was used in this study.

More broadly, the switch from one mRNA isoform to another may alter not just the translational efficiency of the transcript, but localization, stability, or alternative splicing as well. In this regard, transcript toggling driven by developmental switches in promoter usage and the subsequent transcriptional interference from distal gene promoters may serve to alter gene expression in respects other than gene silencing. We posit that the *Adh* example is likely to be one of many cases where developmentally controlled transcriptional interference from ORF-distal promoters can alter genome decoding and cellular function in a manner that has not been anticipated previously.

## Materials and Methods

### Fruit fly stocks, husbandry and larval collection

Fruit flies were raised on standard molasses medium at 25°C. Oregon-R was used as wild type (a generous gift from Don Rio). The *tubGAL4* line was obtained from the Bloomington *Drosophila* Stock Center (ID 5138). All fruit flies in the stock were heterozygous for *tubGAL4* and the balancer *TM3, Sb^1^ Ser^1^*, as the *tubGAL4* chromosome is homozygous-lethal. For experiments requiring adult fruit flies, a mixture of males and females was used. The *Adh^DIST∆^* line was homozygous for the deletion allele. In experiments requiring induction of *Adh^DIST^* in larvae, we crossed homozygous *Adh^UAS^* males to virgin female *tubGAL4/TM3, Sb^1^ Ser^1^* or Oregon-R control fruit flies in collection cages with molasses plates spread with live yeast. After 8 hours, plates were removed, and embryos were allowed to age for 72 hours at 25 °C. The population consisted of predominantly first and second instar larvae. To collect the samples, larvae were washed off the plates using PBS and then washed three times in PBS. In between washes, larvae were left undisturbed to allow settling by gravity. ~2 mL of larvae were aliquotted, flash-frozen in liquid nitrogen, and stored at -80°C for later processing.

### Generation of transgenic fruit flies

We cloned sgRNAs into pCFD4 (Port et al., 2014), which expresses two sgRNAs from U6 snRNA promoters. Two sgRNAs were used to ensure that at least one double-stranded break was formed. The sgRNA plasmid for generating *Adh^DIST∆^* (pUB1041) expressed sgRNAs 5‘-AGTGGGCTTGGTCGCTGTTG-3’ and 5’-TAATATAGAAAAAGCTTTGC-3’. The sgRNA plasmid for generating *Adh^UAS^* (pUB1038) expressed sgRNAs 5‘- CATAACTCGTCCCTGTTAAT-3’ and 5’-ACACATTTGTTAAAAGCATA-3’. The repair templates were cloned into the pGEX-2TK cloning vector (GE Healthcare). To generate the repair template for the *Adh^DIST∆^* allele (pUB1094), two 1-kb homology arms were amplified from Oregon-R genomic DNA, with the *Adh* distal promoter region removed. When used as a repair template donor, this results in the removal of the region spanning -387 to -1376 bp upstream of the proximal isoform transcriptional start site. A similar allele was described previously (Corbin and Maniatis, 1989). The repair template to generate *Adh^UAS^* (pUB1091) contained two 1-kb homology arms amplified from Oregon-R genomic DNA, flanking a *10xUAS-hsp70(core promoter)* construct amplified from pVALIUM20 (Ni et al., 2011). When used as a repair template donor, this results in the insertion of the *10xUAS-hsp70(core promoter)* construct at the -1 position relative to the distal transcriptional start site.

sgRNA plasmids and their corresponding repair templates were injected into *y^1^ w M{nos-Cas9.P}ZH-2A* embryos (Bloomington 54591), which express maternal Cas9, by BestGene Inc. (Chino Hills, CA). The resulting mosaic fruit flies were outcrossed to *w^1118^*, and the F1 progeny were individually crossed to *CyO* or *CyO, twi>GFP* balancer lines prior to being genotyped. Introduction of the desired allele in the genotyped parent was tested by PCR and sequencing. F2 progeny carrying the desired allele and balancer were then crossed *inter se* to generate homozygous animals.

### RNA isolation, cDNA synthesis and Quantitative PCR

Total RNA was isolated using Trizol (Life Technologies) according to a previously described protocol (Bogart and Andrews, 2006). 450 ng of isolated RNA was treated with DNase (TURBO DNA-free kit, Thermo Fisher (MA, USA), and reverse transcribed into cDNA (Superscript III Supermix, Thermo Fisher) according to the manufacturer’s instructions. The RNA levels of specific *Adh* isoforms were quantified using primers specific to Adh^DIST^ and Adh^PROX^ (Table 1), SYBR Green/Rox (ThermoFisher), and the StepOnePlus Real-time PCR system (ThermoFisher). Adh^DIST^ and Adh^PROX^ signals were normalized to *αTUB84B* transcript levels. RT-qPCR for each sample was performed in technical triplicate and the mean Ct value was used for the normalizations. The efficiency value for each oligonucleotide pair was empirically determined and only those pairs that had greater than 90% efficiency were used for the RT-qPCR experiments. The oligonucleotide sequences used for the RT-qPCR experiments are displayed in Table 1.

### RNaseH of Total RNA

To distinguish the size difference between the two *Adh* isoforms, the total RNA of each sample was treated with RNaseH prior to RNA blot analysis. A total of 15 µg Trizol-extracted RNA was added to 1x RNaseH buffer (New England Biolabs, Ipswich, MA). Next, a site-specific DNA oligo (See Table 1 for sequence) was annealed to RNA by heating to 52°C and slowly cooling to 25°C. The RNA-DNA hybrid strands were incubated with 1U RNaseH (New England Biolabs) for 1 hour at 37 °C. RNA was extracted in phenol:chloroform (1:1) and precipitated in isopropanol with 0.3 M sodium acetate overnight at -20°C.

### RNA blotting

RNA blot analysis protocol was performed as described previously (Koster et al., 2014) with minor modifications. 15 ug of total RNA was denatured in a glyoxal/DMSO mix (1M deionized glyoxal, 50% v/v DMSO, 10 mM sodium phosphate (NaPi) buffer pH 6.5–6.8) at 70°C for 10 min. Denatured samples were mixed with loading buffer (10% v/v glycerol, 2 mM NaPi buffer pH 6.5–6.8, 0.4% w/v bromophenol blue) and separated on an agarose gel (1.1-1.5% w/v agarose, 0.01 M NaPi buffer) for 3 hr at 116 V. RNAs were then transferred onto nylon membranes overnight by capillary transfer. rRNA bands were visualized by methylene blue staining. The membranes were blocked in ULTRAhyb Ultrasensitive Hybridization Buffer (Thermo Fisher) for 3 hours before overnight hybridization. Membranes were washed twice in Low Stringency Buffer (2x SSC, 0.1%SDS) and three times in High Stringency Buffer (0.1X SSC, 0.1% SDS). All hybridization and wash steps were done at 42°C. Radioactive probes were synthesized using a Prime-It II Random Primer Labelling Kit (Agilent, Santa Clara, CA). The oligonucleotide sequences of the primers used to generate the *Adh* DNA templates are listed in Table 1.

### Rapid amplification of cDNA ends (5’ RACE) analysis

GeneRacer^TM^ Kit Version L (Life Technologies) was used for full-length, RNA ligase-mediated rapid amplification of 5’ cDNA ends according to manufacturer’s instructions. 2ug of total RNA was isolated, as described above, from L1/L2 larvae and adults. The gene specific primer used is listed in Table 1. The resulting RACE products were analyzed and identified by DNA sequencing.

### H3K36me3 Chromatin immunoprecipitation (ChIP)

Chromatin immunoprecipitation in larval samples was performed as previously described (Alekseyenko et al., 2006) with the following modifications. Chromatin from approximately 2 mL of larval samples was isolated and fixed in 1.0% w/v of formaldehyde for 20 min at room temperature and quenched with 100 mM glycine. Crosslinked chromatin was sonicated 12 times with a 30 seconds ON/30 seconds OFF program using a Bioruptor^®^ Pico (Diagenode, Denville, NJ). A fragment size of ~200 bp was obtained. To preclear the lysate, the samples were incubated in pre-RIPA buffer (10 mM Tris-Cl, pH 8.0, 1 mM EDTA, pH 8.0, 0.1% SDS) containing cOmplete^™^ Protease Inhibitor Cocktail (Roche) and 1mM PMSF with Protein A Dynabeads (Invitrogen) for 2 hours at 4 °C with rotation. After removal of Protein A Dynabeads, pre-cleared lysates were incubated overnight with 4 µg of rabbit anti-mouse IgG (Ab46540, Abcam) and anti-Histone H3K36me3 (Ab9050, Abcam). Simultaneously, a new aliquot of Protein A Dynabeads were blocked in pre-RIPA buffer + 1 µg/µl bovine serum albumin overnight at 4 °C. The immunoprecipitates were then incubated with the pre-blocked Protein A Dynabeads for 4 hours at 4°C. Reverse crosslinked input DNA and immunoprecipitated DNA fragments were amplified with Absolute SYBR green (AB4163/A, Thermo Fisher, Waltham, MA) and quantified with a 7500 Fast Real-Time PCR machine (Thermo Fisher). The efficiency value for each oligonucleotide pair was empirically determined and only those pairs that had greater than 90% efficiency were used for the RT-qPCR experiments. The oligonucleotide sequences of the primers used in RT-qPCR reactions are listed in Table 1.

### Polysome Fractionation and RNA extraction

Whole fruit flies or ovaries were dissected in 1X PBS and were transferred to a microcentrifuge tube on liquid nitrogen. Samples were homogenized on ice in 200 uL cold lysis buffer in presence of cycloheximide. The lysis buffer for cycloheximide samples is as follows: 500 mM KCl, 15 mM Tris pH 7.5, 15 mM MgCl_2,_ 0.5 mM Puromycin, 0.02 U SUPERase-In, 1 cOmplete ULTRA EDTA-free protease inhibitor pill per 50 mL. Samples were centrifuged for 10 minutes at 15,0000 g at 4C. Aqueous phase was transferred to a new pre-chilled microcentrifuge tube, avoiding pellet and wax layer. 10% of the aqueous volume was transferred to a new microcentrifuge tube, with 100 uL TriZol and stored at -80C for mRNA input sample. A 10% sucrose buffer (500 mM KCl; 15 mM Tris-HCl, pH 7.5; 15 mM MgCl2 and 7 uL SuperaseIn) and 50% sucrose buffer (500 mM KCl; 15 mM Tris-HCl, pH 7.5; 15 mM MgCl_2_ and 7 uL SuperaseIn) were used to generate a sucrose gradient of 10% to 40% in a Beckman Coulter 9/16×3.5 PA tube (Cat #331372) SW-41 ultracentrifugation tube. The gradient tube was stoppered and the setting “long sur 10-40%” was used to make the gradient. Gradients were centrifuged at 35,000xg using a SW-41 rotor for 3 hours at 4°C and fractionated on a Brandel flow cell (Model #621140007) at 0.75 mL/min with the sensitivity setting at 0.5 Abs. A volume of 750µl was collected for each fraction. The samples were then pooled as indicated in Figure S1. 5 ng *rcc1*(Xl)-polyA spike RNA was added to each pooled fraction prior to RNA extraction. RNA was extracted from the fractions using standard acid phenol:chloroform extraction as described in (Chan et al., 2018). The RNA pellet was washed with 80% ethanol and then air-dried. After air-drying, the pellet was dissolved in 10 µl of nuclease-free water. The samples were then treated with Turbo DNase prior to cDNA synthesis.

### Data availability

All the reagents generated in this study are available upon request.

## Acknowledgements

We would like to thank Qingqing Wang and Emily Brown for technical help and insight, Don Rio for comments on the manuscript, Jasper Rine for general advice and all members of the Brar and Ünal lab for comments and suggestions. The food for *Drosophila* melanogaster has been prepared and provided by the UC Berkeley Biological Divisional Services Fly Food Facility. This work was supported by funds from the Pew Charitable Trusts (00027344), Damon Runyon Cancer Research Foundation (35-15), National Institute of Health (DP2 AG055946-01) and Glenn Foundation for Medical Research to E.Ü; funds from the Pew Charitable Trusts (00029624), the Alfred P. Sloan Foundation (FG-2016-6229), and the National Institute of Health (DP2 GM-119138) to G.B; funds from the Shurl and Kay Curci Foundation to L.Y.C, a NSF Graduate Research Fellowship to J.C (DGE-1106400) and a NSF Graduate Research Fellowship to E.M.S (DGE 1752814 and DGE 1106400).

## Author contribution

V.J performed the molecular biology experiments including RNA isolation, RNA blotting, cDNA synthesis, RT-qPCR, ChIP and 5’ RACE. J.C, H.V.W, D.H and E.M.S designed and generated the CRISPR/Cas9 transgenic lines, maintained the fruit fly stocks, and performed larval collection. Y.Z assisted in the molecular biology experiments. L.Y.C performed larval polysome fractionation. S.W.W and P.R performed adult polysome fractionation. E.Ü isolated RNA from both polysome fractions. E.Ü, L.Y.C and E.M.S supervised the project. V.J, J.C, H.V.W, E.M.S, L.Y.C G.A.B and E.Ü wrote the manuscript.

## Supplementary Figure Legends

**Supplemental Figure 1.**
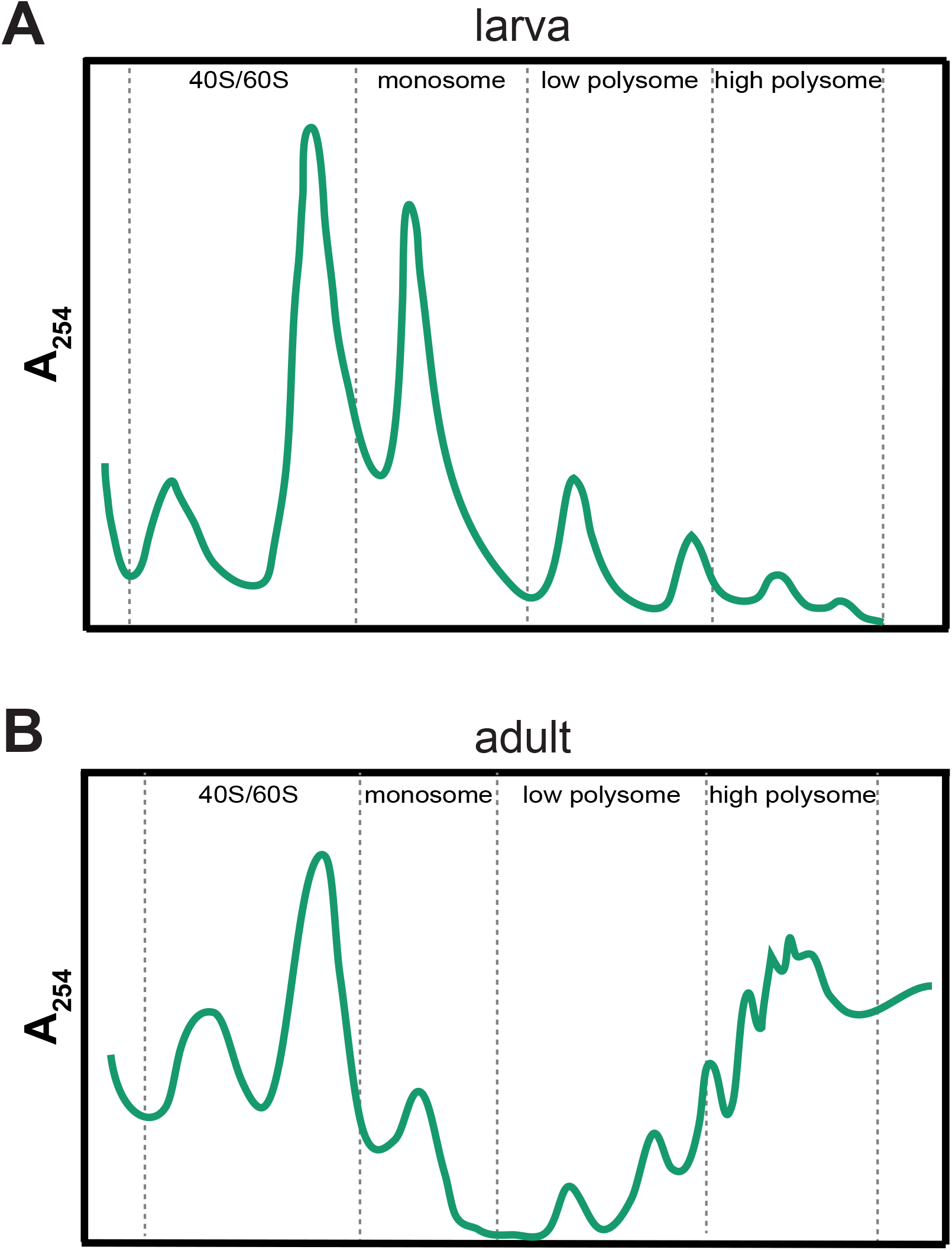
Cytoplasmic lysates from larva (A) and adults (B) were fractionated through sucrose gradients. Pooled fractions used for RNA extraction are highlighted as 40S/60S, monosome, low polysome and high polysome.

**Supplemental Figure 2.**
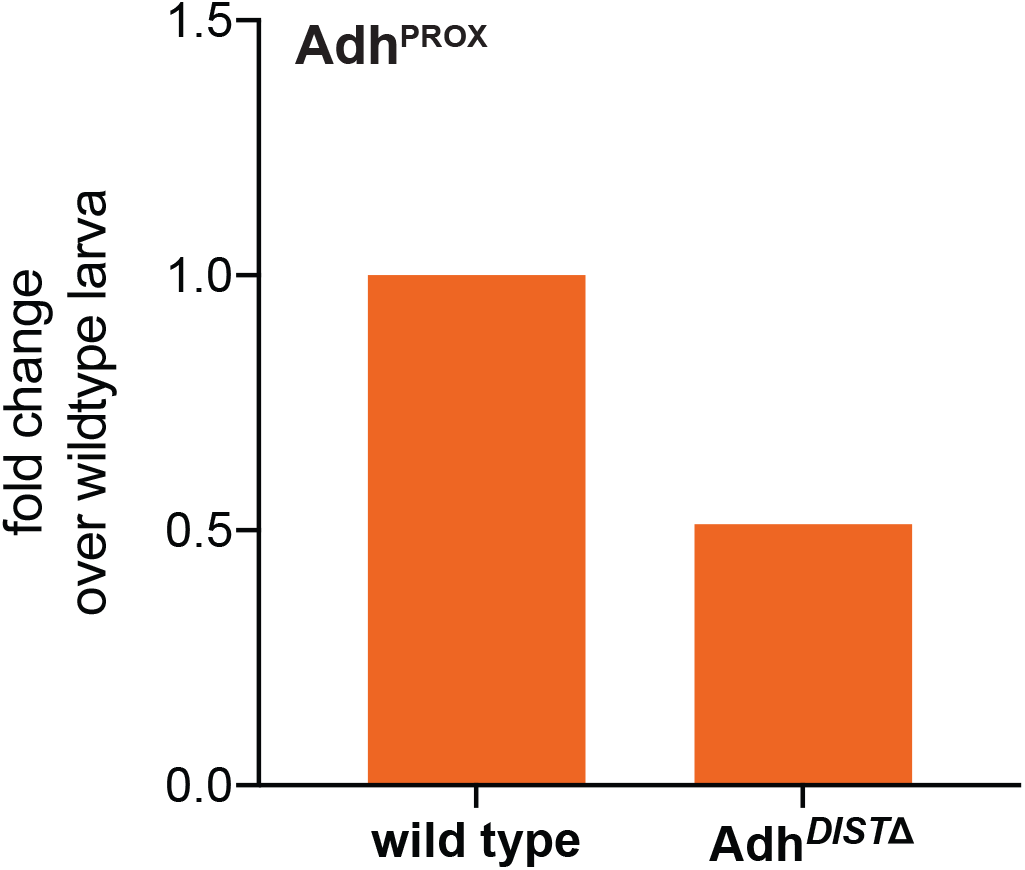
(A) Expression levels of Adh^PROX^ were measured by RT-qPCR using isoform specific primers. Data were normalized to *αTUB84B* and then to wild-type larvae levels to show fold change over wild type (1X).

**Supplemental Figure 3.**
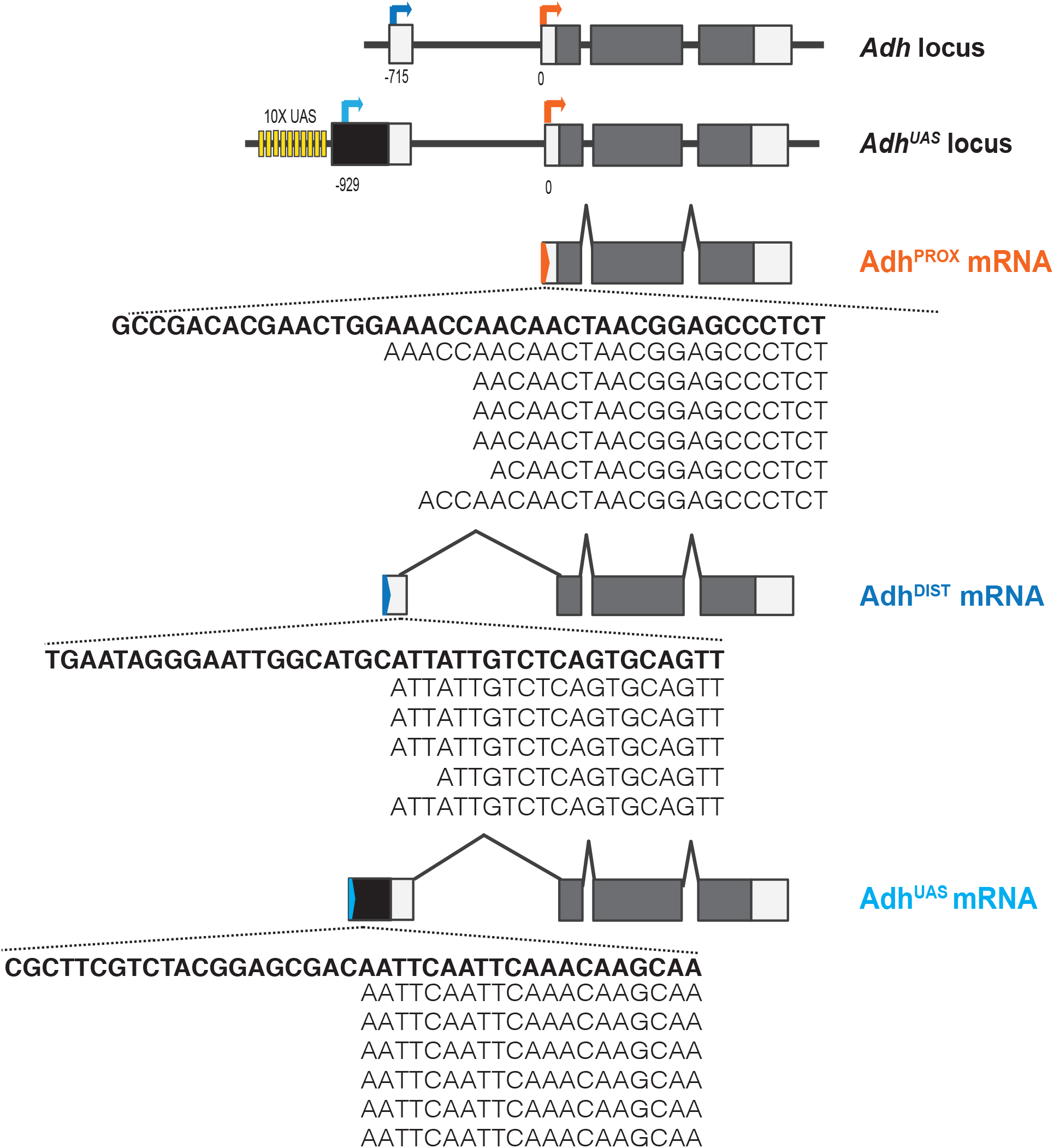
Data from 5’ Rapid amplification of cDNA ends (5’ RACE) analysis for Adh^PROX^, Adh^DIST^, and Adh^UAS^. At the top are diagrams of the wild-type *Adh* locus and the *Adh^UAS^* locus with the GAL4/UAS induction system. At the *Adh^UAS^* locus, 10 consecutive GAL4 binding sites (UAS) (shown in yellow) are placed immediately upstream of the Adh^DIS*T*^ TSS followed by the minimal *hsp70* promoter (black). Coding (grey) and non-coding (white) exons are shown. Arrows represent TSSs of Adh^PROX^ (orange), Adh^DIST^ (blue) and Adh^UAS^ (light blue). Numbers below the *Adh* locus refer to distance in base pairs (bp) from the Adh^PROX^ TSS. Diagrams for each isoform’s mRNA transcript are shown above the sequence results. Genomic DNA sequence is bolded with aligned transcript sequences listed below.

